# Limits of marker information transfer in cross-panel cytometry imputation

**DOI:** 10.64898/2026.10.07.757120

**Authors:** Jonghyun Lee, Akira Nair, Chang-Uk Jeong, Matei Ionita, Jaehyun Joo, Matthew E. Lee, Lina Takemaru, Yonghyun Nam, Jaesik Kim, Byounghan Lee, Michelle L. McKeague, Shwetank Verma, Ayman Rezk, David Smith, Sokratis A. Apostolidis, Amy E. Baxter, Martin P. Carroll, Allison R. Greenplate, E. John Wherry, Dokyoon Kim

## Abstract

Marker imputation can bridge cytometry datasets acquired with different panels and platforms. Its validity depends on whether common markers retain information about unmeasured target markers and whether those relationships transfer across assay contexts. We evaluated these limits in two paired flow cytometry–CyTOF cohorts comprising 134 patients and 257 specimens. A patient-held-out framework distinguished within-platform same-cell reconstruction from cross-platform distribution translation without cell correspondence. We compared conventional machine-learning and deep generative estimators with median imputation and used permutation controls to estimate the contributions of patient matching and source-exclusive markers. Evaluation covered marker values, specimen distributions, phenotype utility, and directional predictive relationships among markers. Within platforms, exact k-nearest neighbors reduced mean absolute error by 12.8–35.0% relative to median imputation. However, the median-based reconstructions were contracted, with a median ratio of predicted to observed interquartile range below 1 in every context and evaluable population. Patient matching improved cross-platform distribution recovery, whereas adding source-exclusive markers or higher-capacity estimators did not consistently produce net gains. In the independently annotated acute myeloid leukemia (AML) cohort, reconstructed markers largely preserved broad-lineage classification but did not recover fine-phenotype information available through direct measurement. In AML–CyTOF, the measured full panel outperformed the reconstructed representation by 10.25 percentage points for fine phenotypes. Predictive relationships among markers were context-specific. Most targets were better explained by several markers than by one. Marker selection based on these relationships showed a trade-off between panel size and pairwise coverage. Paired samples and imputation can therefore connect selected marker distributions across panels but should not be treated as substitutes for direct measurement. Cross-panel cytometry should be evaluated with patient-held-out tests matched to the resolution of available ground truth and the intended downstream use.

**Key Points:**

- Within-platform masking and cross-platform patient-matched translation answer different questions and should be evaluated at the resolution of their available ground truth.
- Patient matching improved cross-platform distribution recovery, whereas adding source-exclusive markers or higher-capacity estimators did not consistently produce net gains.
- Reconstructed markers largely preserved broad-lineage classification, but in AML–CyTOF they did not recover fine phenotypes such as *γδ* and double-negative T cells, which are separated by a marker absent from the common panel.
- Predictive relationships among markers were context-specific and multi-source, and graph-guided panels with high pairwise coverage did not necessarily preserve phenotype utility.
- Cross-panel imputation should be validated with patient-held-out tests matched to the intended downstream use, and markers central to decisions should be measured directly.

## Introduction

Cytometry measures the expression of multiple protein markers simultaneously in tens of thousands to millions of single cells, capturing immune composition and cellular state at high resolution. Marker panels and measurement platforms, however, vary with study objectives and assay design, so related biology is often observed only partially and in different marker spaces and measurement scales. These differences impede direct comparison and integration across studies(29).

Immunologists have traditionally defined cell populations by manual gating and interpreted data through population abundance and marker-expression summaries. This approach is interpretable but labor-intensive and sensitive to the analyst and gating strategy. It also reduces continuous cell-state variation to discrete populations and summary statistics. To mitigate these limitations, machine-learning and deep-learning models use cell-level expression directly to automate population annotation and sample-level prediction while retaining high-dimensional biological structure(3; 21; 29; 20; 4; 34; 14).

Two related directions have recently emerged in computational cytometry. First, self-supervised and large-scale pretrained models learn representations that can be reused across datasets and downstream tasks. cyMAE(14), ImmuneFM(8), CytoBERT(28), and scpFormer(35) learn cell-level representations from single-cell marker-expression patterns, with recent work extending to heterogeneous or variable panels. MAESTRO(17) and EventHorizon(22), by contrast, aggregate entire cell sets into sample- or specimen-level representations.

Second, several methods explicitly connect partially overlapping panels to impute unmeasured markers. Infinity Flow(5) predicts protein expression from backbone markers. Multitube profiling(25), CyTOFmerge(1), and CytoBackBone(19) transfer information between similar cells in common-marker space. Data-fusion approaches combine complementary panels for sample-level prediction(31) Within common-marker clusters, cyCombine(26) restores missing-marker distributions. CytoVI(13) and UVAE(27) use shared latent spaces to align panels and reconstruct unmeasured markers.

Representation learning and explicit imputation produce different outputs but depend on the same premise: common markers must contain sufficient information about target markers and cellular state, and those relationships must transfer across panels and biological contexts. Representation learning seeks to capture this shared structure, whereas imputation asks whether unmeasured values can be recovered from it. The question is therefore how much target-marker information remains in common markers and whether that information persists in a new dataset or platform.

Many studies evaluate reconstruction by computationally masking markers or splitting a native full-panel dataset into pseudo-panels(1; 14; 27). These experiments provide same-cell ground truth and quantify marker predictability, but source and target measurements arise from the same acquisition context. A previous critical assessment further showed that plausible imputed distributions can coexist with errors in cell-level expression and gating(23). In a true cross-platform setting, even common markers differ in scale and noise structure across instruments and assays. Platform and panel may also be confounded with disease or cohort(26; 27). Paired-patient measurements connect biological context but do not establish correspondence between individual cells from different aliquots. Within-platform cell-level reconstruction must therefore be distinguished from cross-platform translation evaluated at the sample or population distribution level.

Panel composition determines the biological information accessible to a model. Imputation, in particular, depends on the information carried by the common markers(1; 26; 23). If target variation is not shared with common markers, or if marker relationships do not persist across contexts, increasing model complexity or training scale cannot restore that information. Conversely, quantifying stable marker relationships may support panel design that captures more information under a constrained measurement budget.

We evaluated the limits of within-platform and cross-platform marker recovery in four cytometry contexts from two paired-platform cohorts using a patient-held-out framework (Figure 1). We asked three questions: what target-marker information is accessible from common markers and paired samples, whether reconstructed markers retain downstream phenotype utility, and whether predictive-information structure is sufficiently stable to guide marker prioritization. Within-platform masking quantified cell-level and distributional recoverability. Across platforms, a patient-matched bridge evaluated sample- and population-level distribution translation without assuming individual-cell correspondence. We compared median imputation, conventional machine-learning estimators, and deep generative models; used permutation controls to estimate the contributions of patient matching and source-exclusive markers; and evaluated phenotype utility and directional predictive-information structure. This design separates the information that panels and pairing make available from how well estimators use it. It also identifies where direct measurement remains necessary.

**Figure 1.**
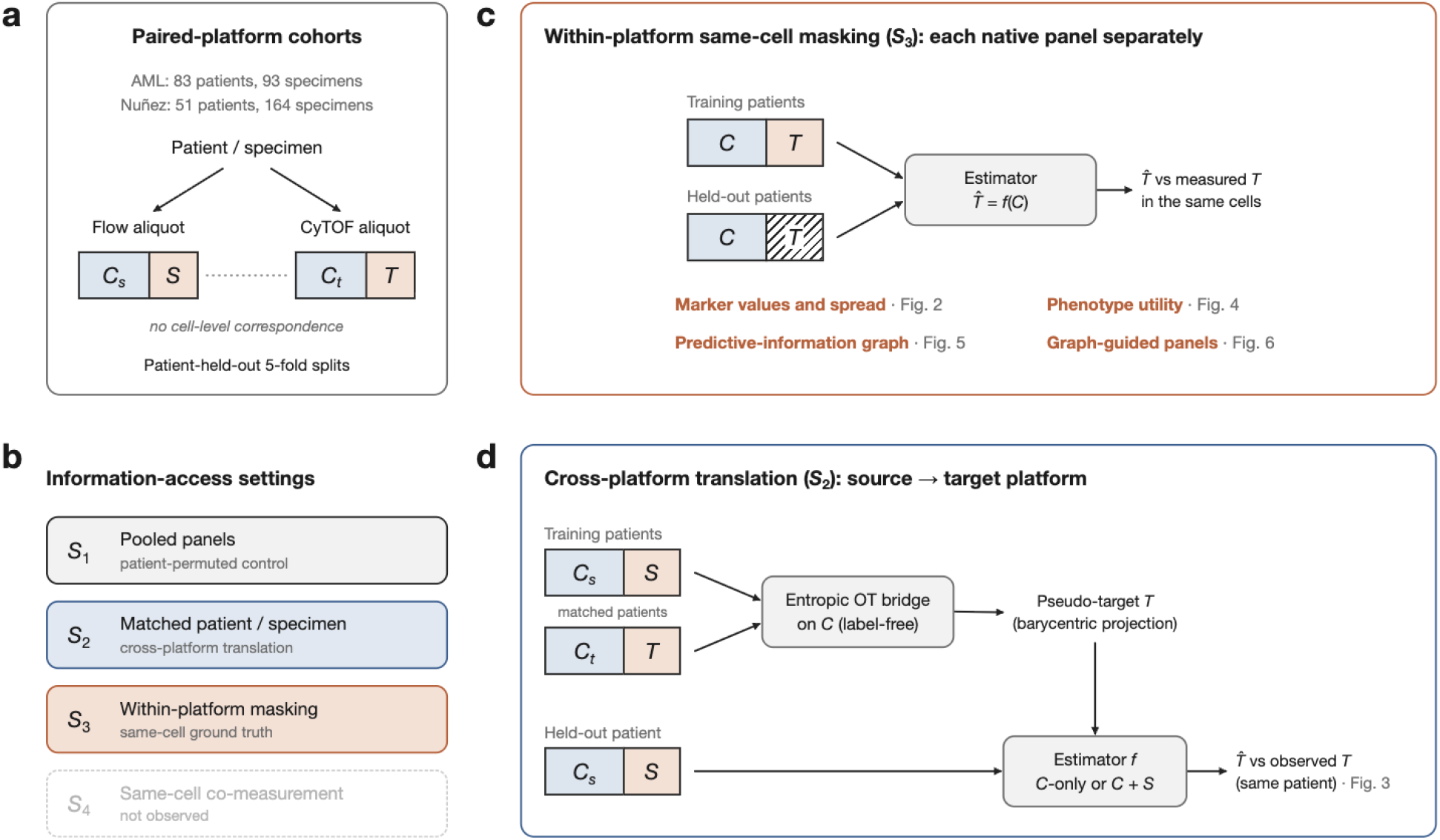
Study design. (a) Each patient or specimen was measured on separate flow cytometry and CyTOF aliquots, sharing common markers (*C*_*s*_, *C*_*t*_) but no cell-level correspondence. All evaluations held out entire patients. (b) Information-access settings. (c) Within-platform masking (*S*_3_) masked native exclusive markers in held-out patients and compared reconstructions with measured values in the same cells. This setting supported marker-level, phenotype, predictive-information, and panel-selection analyses. (d) Cross-platform translation (*S*_2_) used a label-free optimal-transport bridge on common markers to build target-exclusive pseudo-targets from matched training patients and evaluated reconstructed target-marker distributions against the held-out patient’s observed target distributions. Patient-permuted and within-specimen permuted (*S*_perm_) controls estimated the contributions of patient matching and source-exclusive inputs.

## Materials and Methods

### Problem Definition

.We considered paired settings in which the same patient or specimen was measured on different cytometry platforms. For one translation direction, let *C* denote markers measured on both the source and target panels, *S* source-exclusive markers, and *T* target-exclusive markers. The source therefore measures (*C*_*s*_, *S*) and the target measures (*C*_*t*_, *T* ). Although *C*_*s*_ and *C*_*t*_ refer to the same biological targets, we did not assume equal values because they were obtained from different instruments and aliquots. Reversing the translation direction also reverses the roles of *S* and *T* .

Patient matching links sample context, including disease state, time point, and cell-population composition, but does not link individual cells. Cross-platform data therefore directly identify only same-cell marker relationships within each platform and distributions across matched samples. We used this distinction to separate two questions: at what resolution an unmeasured marker can be recovered, and whether its reconstruction adds phenotype information.

### Datasets and Processing

We analyzed two paired-platform cohorts.

- **I3H AML dataset:** Ninety-three specimens from 83 patients with acute myeloid leukemia (AML) were measured by spectral flow cytometry and CyTOF. The spectral-flow and CyTOF panels contained 26 and 44 markers, respectively, including 19 common markers. The platforms shared patient/specimen keys but used separate aliquots and thus had no cell-level correspondence.
- **Nuñez dataset:** We used 164 specimens or repeat visits from 51 patients available in the public release of a paired cohort(24). The FACSymphony and CyTOF panels contained 26 and 37 markers, respectively, including 20 common markers. Repeat visits from the same patient were assigned to the same patient-level split.

We refer to the spectral-flow and FACSymphony contexts as AML–flow and Nuñez–flow, respectively, and to the two mass-cytometry contexts as AML–CyTOF and Nuñez–CyTOF. Cross-platform analyses are identified by their explicit source-to-target translation direction.

The I3H AML data were processed with the institutional quality-control pipeline and annotated with immunologist-defined manual-gating labels. For the Nuñez cohort, we processed the publicly released paired FCS files and did not use author-provided percell phenotype annotations. We derived six analysis-specific coarse lineage strata (CD4 T, CD8 T, other T, B, NK, and other) from CD3, CD4, CD8, CD19, and CD56 using specimen-specific positivity thresholds. These strata were used for stratified analyses and were not treated as investigator-provided manual-gating labels. Marker names were harmonized to canonical names. The primary analysis fixed the 19 AML and 20 Nuñez native common markers as input *C* and treated the remaining markers in each direction as *S* or *T* .

All analyses used five outer cross-validation folds grouped by patient. Marker-specific distances were standardized using scales estimated with equal total weight per outer-training patient. We used the interquartile range (IQR) for most markers. For nonconstant markers with zero IQR, we fell back to the median absolute deviation and then to the winsorized root-mean-square. Truly constant markers were assigned a scale of 1. This procedure prevented nearly invariant markers from dominating distance calculations.

### Information-Access Settings

Recoverable information depends on the linkage available between platforms.

*S*_1_ **Pooled panels:** Only cohort-level distributions are available, without patient matching.

*S*_2_ **Matched patient/specimen:** Sample- and population-level translation can be evaluated without cell pairs.

*S*_3_ **Within-platform masking:** Same-cell truth within a native panel permits cell-level reconstruction evaluation.

*S*_4_ **Same-cell co-measurement:** Cross-platform same-cell recovery could be evaluated directly, but this setting was not observed in our study.

*S*_1_ was a baseline without the contribution of patient matching. We approximated it with a patient-permuted control. The primary analyses focused on *S*_2_ and *S*_3_.

### Study Design and Evaluation Framework

#### Information access and recoverability

##### Within-platform masking

Because cross-platform data do not provide same-cell truth, we first masked markers within each native full panel. We used *C* as input, masked native exclusive markers *T*, and evaluated predictions against values originally measured in the same cells. This experiment quantified both cell-level error and distribution recovery in held-out specimens. The four contexts were evaluated independently.

##### Cross-platform patient-matched translation

In the cell-unpaired *S*_2_ setting, we transformed outer-training source and target *C* into patient-balanced percentile space. We then used label-free entropic optimal transport with barycentric projection to construct target-exclusive pseudo-targets *T* (7). Regularized optimal transport has previously been used to estimate cell-population proportions in an unlabeled cytometry sample from a manually gated one(10). Cell-type and endpoint labels were withheld from both the bridge and the predictor.

For each translation direction, we compared *C*-only, *C* + *S*, and *C* + *S*_perm_ inputs. The *S*_perm_ control permuted source-exclusive marker vectors among cells within each specimen. It preserved dimensionality, marginal distributions, and joint structure among exclusive markers while removing the link between each vector and the corresponding cell’s *C*. A separate patient-permuted control mapped each source patient to a different target patient by derangement, removing sample-level information supplied by patient matching.

##### Estimator comparison

To test whether estimator complexity consistently improved recovery, we compared median imputation, conventional machine-learning models, and cytometry-specific deep generative models. Median imputation was a null reference that did not use input markers. This analysis focused on marker reconstruction and distribution fidelity. Downstream phenotype utility was evaluated separately.

#### Downstream phenotype utility

We evaluated phenotype utility using cohort-specific label endpoints: immunologist-defined manual-gating labels for AML and analysis-defined coarse lineage strata for Nuñez. We compared the macro-averaged area under the precision-recall curve (macro-AUPRC) among *C*, 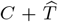, and the measured full representation (*C* + *T* ). Fine-label and one-vs-rest analyses were restricted to AML.

#### Predictive-information structure and graph-guided marker selection

##### Directional predictive-information graph

We estimated a directional predictive-information graph from same-cell measurements in each native panel, extending conditional information measures previously used to quantify marker relationships in mass cytometry(15). An edge *M → T* measured the reduction in held-out uncertainty about target *T* when source marker *M* was added to a prediction based on the cell label alone. We subtracted a null obtained by permuting markers within patient and label. This step reduced apparent relationships attributable only to patient-specific composition or marker abundance.

Across cytometry contexts, we evaluated preservation of common-marker edge magnitudes and node-coverage ranks. Within each outer-training fold, we also selected the two leading sources for each target and classified their held-out relationship as complementary, redundant, or single-dominant according to whether the second source added information. Detailed estimation and inference procedures are provided in the Supplementary Methods.

##### Graph-guided marker selection

Within each context, markers were added greedily to maximize pairwise coverage of targets not yet measured. Pairwise coverage credited each target with its strongest single selected source. At every panel size *K*, we compared coverage with random panels of the same size. We then identified the smallest *K* that first reached 95% of an oracle in which any other marker could serve as that source. We call this panel size *K*_95_. Because native panel size *D* differed across contexts, panel sizes were compared as *K/D*. The *K*_95_ panel was reselected within each outer-training fold and evaluated by held-out k-nearest-neighbor (kNN) reconstruction and cohort-specific phenotype macro-AUPRC (AML fine labels and Nuñez coarse strata). Because *K* was chosen post hoc from the aggregate graph, this analysis was interpreted as an exploratory sensitivity analysis. The analysis code calculates the greedy selection path, *K*_95_, and random-panel reference from an edge table. An interactive exploration of predictive-information graphs and graph-guided marker selection is shown in Supplementary Figure S1.

### Baseline Models

Models were divided into inductive and transductive tracks according to whether the held-out query distribution was used during fitting. Inductive models were trained only on outer-training references and then predicted query cells independently. Transductive models did not observe query targets or labels but used the common-marker distribution of the held-out query during fitting. They were therefore reported as a separate track.

#### Median imputation

Median imputation assigned the patient-balanced outer-training target median to every query cell. In cross-platform translation, this target was the bridged pseudo-target. Because median imputation did not use cell-to-cell variation in the input markers, it defined the 0% reference for relative mean absolute error (MAE) reduction.

#### Inductive models

Ridge regression learned a linear mapping from *C* to each target. The kNN estimator identified the 50 nearest outer-training cells in standardized *C* and returned the marker-wise median target value among the neighbors, following CyTOFmerge(1). Within-platform references were capped at 5,000 cells per patient and 50,000 cells overall. Cross-platform references were the patient-balanced 50,000-cell bridge bank. Gradient-boosted trees fit a nonlinear ensemble for each target, and the multilayer perceptron (MLP) learned a nonlinear mapping with two hidden layers. MLP epochs were selected by inner patient-level validation. All models were evaluated on the same held-out patients.

#### Transductive models

CytoVI is a probabilistic generative model that aligns partially observed panels in a shared latent space and decodes unmeasured markers(13). UVAE combines panel-specific variational autoencoders with alignment constraints to connect heterogeneous panels in a common latent representation(27). Both models used outer-training full-panel references together with held-out query *C* but did not use query targets or cell labels. Every input condition and model used the same entropic optimal-transport bridge to construct pseudo-targets, so the bridge was not itself a comparator. The cross-platform comparison therefore evaluated estimators after a shared bridge rather than each method’s native panel-integration pipeline.

The all-model comparison used identical patient splits, *C*-only inputs, and target markers. Models were ranked by normalized Wasserstein distance (WD) within each translation direction, and ranks were averaged with equal weight across the four directions. Detailed population- and marker-level analyses used exact kNN as a fixed reference because it did not use the query distribution during fitting.

### Evaluation Metrics

Evaluation was divided into marker values, sample distributions, phenotype classification, and predictive-information structure. Detailed formulas and secondary metrics are provided in the Supplementary Methods.

#### Reconstruction fidelity

In *S*_3_, where same-cell truth was available, relative MAE reduction versus median imputation measured recovery of marker values, and Spearman correlation measured preservation of cell ordering. Positive MAE reduction indicated lower error than median imputation. Distribution fidelity was evaluated with WD, defined as the 1-Wasserstein distance for each held-out specimen and target marker divided by the outer-training target scale, and with IQR retention, the predicted IQR divided by the observed IQR. Lower WD and IQR retention closer to 1 indicated better agreement with the observed distribution. WD improvement for model comparison was calculated as 100(1 *−* WD_model_*/*WD_median_). Because *S*_2_ had no cell pairs, it was evaluated only at the sample and population distribution levels.

#### Cross-platform contrasts

For pairing comparisons, median-imputation WD improvement was set to 0%, and patient-permuted and patient-matched translation were expressed relative to that reference. Larger values therefore indicated better distribution recovery than median imputation. The pairing contribution was patient-matched minus patient-permuted improvement, in percentage points. Source-exclusive input was assessed by direct comparisons among *C*-only, *C* + *S*_perm_, and *C* + *S*.

#### Phenotype classification

AML major phenotype labels were broad manual-gating classes harmonized across platforms, whereas AML fine labels were dataset-native manual-gating classes. The Nuñez cohort had no independent fine-label endpoint, so only the six analysis-defined coarse lineage strata were used. For the native-panel population summary, Blast, B cell, CD4 T, CD8 T, monocyte, and NK cell were grouped as major populations, and Other T and Other were grouped as minor/other. Classifiers were refit separately for *C*, 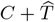, and measured full (*C* + *T* ) representations, and macro-AUPRC was compared. The primary contrast was the AUPRC of each incomplete representation minus that of measured full. Reconstruction gain was 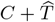 minus *C*-only AUPRC.

#### Predictive-information graph

The magnitude of edge *M → T* was the fraction of held-out target uncertainty reduced by adding *M* . A stable edge met patient-level uncertainty, multiple-testing, and 1% effect-size criteria. Preservation across contexts was evaluated primarily by Spearman correlation of common-edge magnitudes, with stable-edge overlap and node-coverage rank correlation as secondary measures. We prespecified *ρ ≥* 0.60 as the threshold for practical transfer.

#### Aggregation and inference

Within each patient, results were first averaged across specimens and, where repeated, across seeds. Patients were then weighted equally. We call this summary the patient-first mean. For the all-model comparison, overall error was averaged across target markers within each patient before summarizing across patients. Statistical uncertainty was estimated with 2,000 bootstrap replicates using patients as the resampling unit. Multiple testing was controlled with the Benjamini–Hochberg procedure. Detailed permutation procedures are provided in the Supplementary Methods.

## Results

We evaluated four cytometry contexts from two cohorts (AML–flow, AML–CyTOF, Nuñez–flow, and Nuñez–CyTOF). Every comparison was performed in held-out patients. Results are organized around three themes: information access and recoverability, downstream phenotype utility, and predictive-information structure and graph-guided marker selection.

### Information Access Defines the Limits of Marker Recovery

#### Common markers support partial same-cell recovery

With exact kNN, patient-first mean MAE was 12.8–35.0% lower than with median imputation (Table 1). Improvements across the 55 target markers in the four contexts nevertheless ranged widely, from *−*3.6 to 52.0% (Figure 2). Context-specific marker-median Spearman correlations ranged from 0.334 to 0.593, indicating partial preservation of cell ordering. Median IQR retention ranged from 0.346 to 0.599 and was below 1 in every context and every evaluable cell type (Supplementary Table S1). Thus, *C* contained information predictive of unmeasured markers, but accuracy and distribution recovery varied by marker. The median-based reconstructions were also systematically contracted.

**Table 1.**
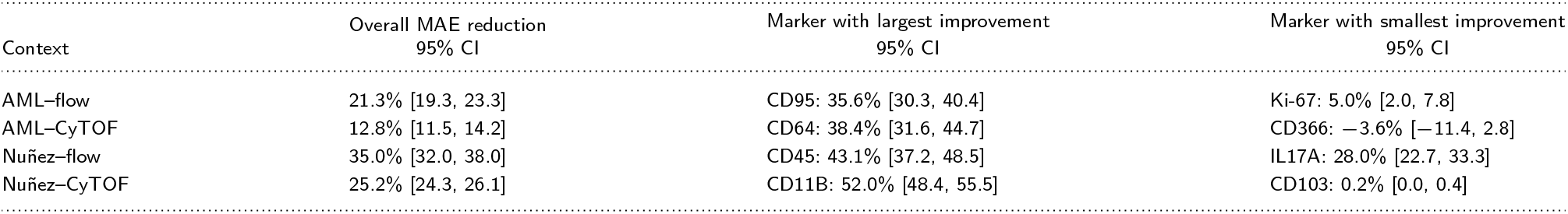
Exact-kNN marker reconstruction in native panels. Values are patient-first MAE reductions relative to median imputation with 95% patient-bootstrap confidence intervals. Positive values indicate lower error; markers with the largest and smallest improvements are descriptive extrema.

**Figure 2.**
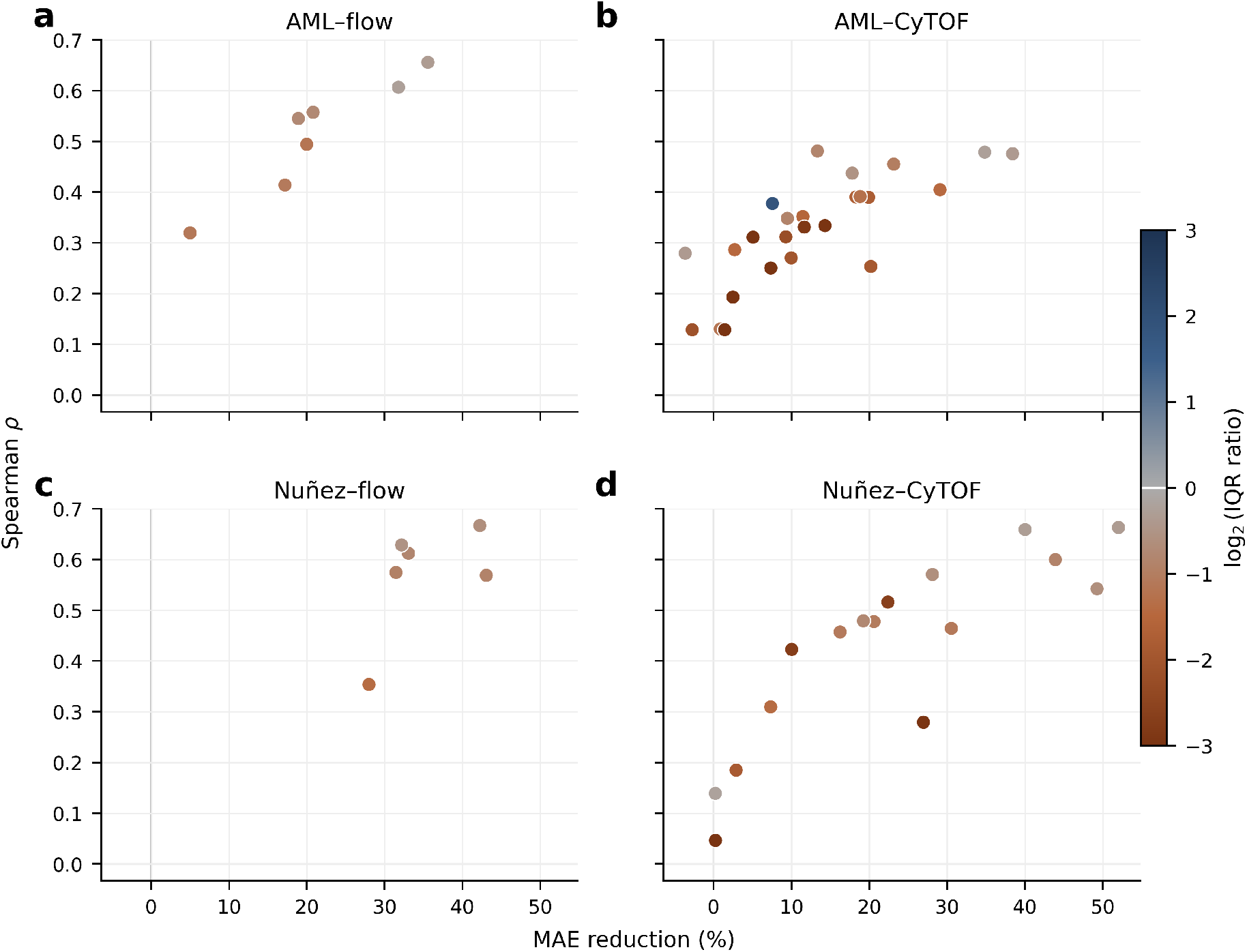
Marker-level fidelity of exact-kNN reconstruction in within-platform masking: (a) AML–flow, (b) AML–CyTOF, (c) Nuñez–flow, and (d) Nuñez– CyTOF. Each point represents one target marker. The x-axis shows MAE reduction relative to median imputation, and the y-axis shows same-cell Spearman correlation between observed and reconstructed values. Color represents the log_2_-transformed patient-first mean ratio of predicted to observed IQR. A value of 0 indicates equal spread, negative values indicate contraction, and positive values indicate expansion. The color range was limited to *±*3 for visualization. Specimens were first averaged within patient, and patients were then weighted equally.

#### Patient matching supports distribution-level translation

With median imputation set to 0%, WD improvement ranged from 9.4 to 14.9% for patient-permuted translation and from 11.7 to 37.7% for patient-matched translation (Table 2). The pairing contribution ranged from 2.4 to 25.9 percentage points across the four translation directions, with confidence intervals above zero in every direction. Within major populations, however, the advantage of patient-matched over patient-permuted translation was less consistent and was absent in every population for Nuñez–flow*→*Nuñez–CyTOF (Supplementary Table S4). Part of the pairing contribution may therefore reflect matched population composition.

**Table 2.** Cross-platform distribution recovery by patient pairing. Values are WD improvements over median imputation (0% reference), averaged across target markers; larger values are better. All conditions used *C*-only exact-kNN prediction. Pairing contribution is patient-matched minus patient-permuted improvement, in percentage points, with patient-bootstrap 95% CIs. Figure 3a reports the ratio of direction-level mean WDs instead.

| Translation context | Patient-permuted | Patient-matched | Pairing contribution [95% CI] |
| --- | --- | --- | --- |
| AML-flow→AML-CyTOF | 14.6% | 27.3% | +12.6 pp [11.4, 13.9] |
| AML-CyTOF→AML-flow | 14.9% | 26.7% | +11.8 pp [10.4, 13.1] |
| Nuñez-flow→Nuñez-CyTOF | 9.4% | 11.7% | +2.4 pp [1.8, 3.1] |
| Nuñez-CyTOF→Nuñez-flow | 11.9% | 37.7% | +25.9 pp [19.2, 31.6] |

In a separate input contrast, *C* + *S* produced lower WD than *C* + *S*_perm_ in all four directions (Supplementary Table S3). Preserving the cell-level association between *S* and *C* therefore improved translation relative to within-specimen shuffling. However, *C* +*S* did not improve on *C*-only in any direction and had higher WD in both CyTOF*→*flow directions.

#### Higher-capacity estimators do not consistently improve recovery

We compared seven models, including median imputation, using identical patient splits, *C*-only inputs, and the shared pseudo-target bridge. All six estimators improved patient-first target-marker mean WD over median imputation, but the best-performing model differed by translation direction (Figure 3a). Exact kNN achieved the best mean rank across the four directions (2.00), whereas UVAE, the best-ranked deep generative model, had a mean rank of 3.25.

**Figure 3.**
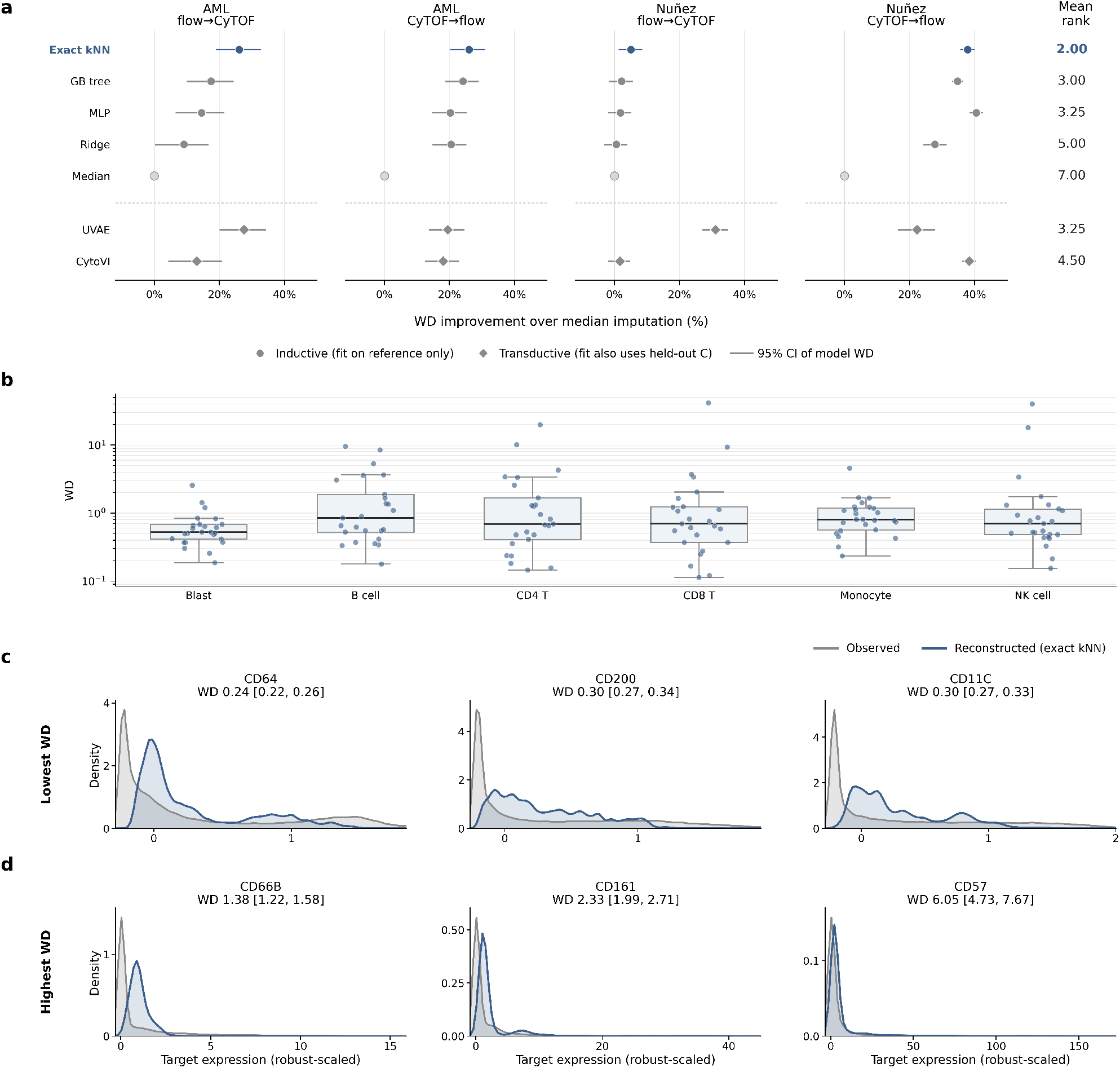
Cross-platform distribution recovery. (a) Patient-first improvement in imputed-distribution WD over median imputation for seven models and their mean rank across four translation directions. Circles denote inductive models and diamonds transductive models. Exact kNN is highlighted in blue. GB tree, gradient-boosted trees. Whiskers show the patient-bootstrap 95% CI of each model’s WD relative to the median-imputation estimate and are not paired contrasts. Unlike the native-panel model sensitivity analysis, all models here shared one pipeline and bridge and were therefore ranked together. (b) Target-marker WD by strict major population for exact-kNN AML–flow*→*AML–CyTOF translation (log scale). Each point represents one marker. Boxes show the median and interquartile range across markers. (c–d) Patient-balanced whole-cell densities for the three target markers with the lowest (c) and highest (d) patient-first WD in exact-kNN AML–flow*→*AML–CyTOF translation. Titles give WD [95% CI]. Gray denotes observed targets and blue denotes exact-kNN reconstructions. Expression is in robust-scaled units, so high WD for sparse markers such as CD57 reflects long tails relative to a small scale. Cross-platform density recovery does not imply same-cell correspondence.

Detailed analysis of AML–flow*→*AML–CyTOF with exact kNN also showed wide variation in target-marker WD within populations (Figure 3b). In CD8 T cells, WD was 0.121 [95% CI 0.117, 0.125] for CD64 but 41.534 [37.710, 45.512] for CD57. Patient-balanced densities for the three markers with the lowest and highest WD likewise differed in their agreement with the observed distributions (Figure 3c,d). Aggregate model ranks therefore obscured marker- and population-specific failures.

In the separate native-panel model sensitivity analysis, kNN was the only estimator with lower MAE than median imputation in all four contexts (Supplementary Table S5). Gradient-boosted trees and the MLP matched or exceeded kNN in the two flow contexts but had higher MAE than median imputation in AML–CyTOF (improvements of *−*4.5% and *−*1.6%, respectively). Inductive and transductive models were evaluated in separate prespecified pipelines. Within either model class, greater complexity did not consistently improve reconstruction of unmeasured markers across contexts.

### Broad-Lineage Performance Was Largely Preserved, Whereas AML Fine-Phenotype Information Was Not

We next asked whether reconstruction preserved phenotype information in within-platform masking (*S*_3_). Phenotype classification with *C*, 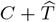, and measured full (*C* + *T* ) representations differed by context and label resolution (Figure 4). After representation-specific refitting, major-label AUPRC minus that of measured full ranged from *−*3.0 to 0.5 percentage points for *C* and from *−*3.1 to 1.3 points for 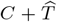. For overall fine-label AUPRC, 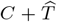 minus *C* was 0.02 points [95% CI *−*0.16, 0.24] in AML–flow and *−*0.20 points [*−*0.45, 0.07] in AML–CyTOF. By contrast, measured full outperformed 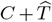 by 0.77 and 10.25 points, respectively. In AML– CyTOF, the gaps for T cell *γδ* and T cell DN (double-negative) were 33.98 and 31.81 points. TCR*γδ* separates these two populations but was not a common marker, so only measured full contained it.

**Figure 4.**
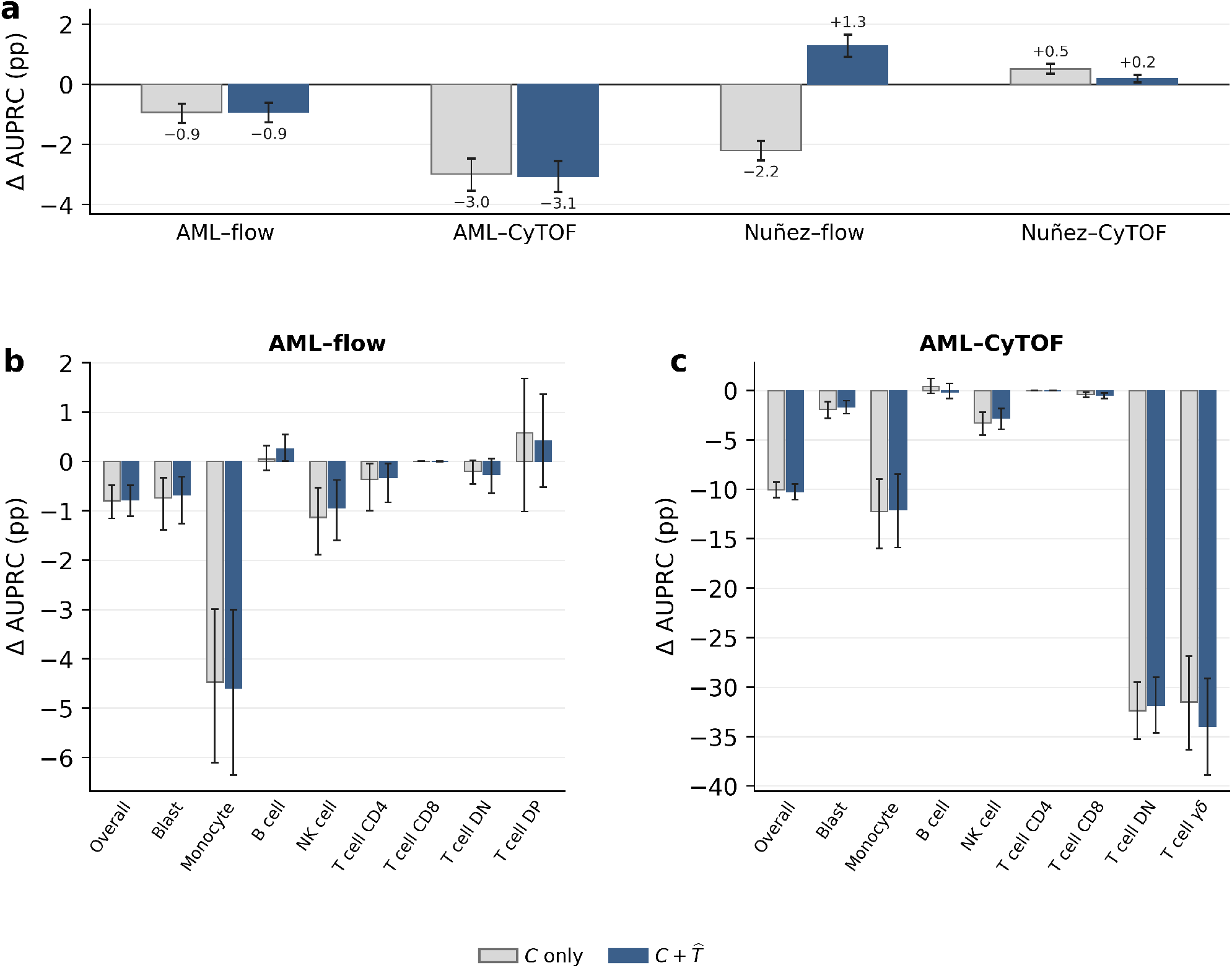
Phenotype classification with reconstructed markers in within-platform masking (*S*_3_). (a) Differences in broad-label macro-AUPRC across the four contexts. AML labels were harmonized manual-gating classes, whereas Nuñez labels were analysis-defined coarse lineage strata derived from markers in *C*. The Nuñez comparison is therefore an internal rule-consistency analysis. (b–c) Differences in overall fine-label and native fine-label one-vs-rest AUPRC in AML–flow and AML–CyTOF. Every value is *C*-only or 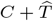 minus measured full (*C* + *T* ), in percentage points. A value of 0 therefore indicates performance equal to measured full, and negative values indicate lower performance. Classifiers were refit for each representation. Error bars are patient-bootstrap 95% confidence intervals. Y-axis ranges differ by context so that changes within each context are visible. DN, double-negative; DP, double-positive.

Because the Nuñez strata were constructed from CD3, CD4, CD8, CD19, and CD56, all included in *C*, their broad-label analysis was interpreted as an internal rule-consistency analysis rather than independent phenotype validation. In this setting, 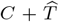 slightly exceeded measured full in Nuñez–flow (1.3 points). Thus, the measured full panel is not a strict upper bound when the endpoint is defined from *C*.

Applying the measured-full classifier directly to 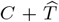 lowered AML–CyTOF fine-label AUPRC below *C*-only. Refitting recovered 4.5 points [95% CI 3.7, 5.3] (Supplementary Table S6). Most of the loss relative to *C*-only under a fixed classifier therefore reflected classifier–representation shift, although a gap to measured full remained after refitting.

### Predictive Information Is Context-Specific and Combinatorial

#### Predictive relationships are weakly preserved across contexts

Same-cell predictive-information graphs differed substantially among contexts (Figure 5; Supplementary Table S8). Stable edges comprised 11.0–78.9% of tested edges, and strong edges that reduced target uncertainty by at least 5% comprised 0.4–10.7%. AML–CyTOF had the sparsest graph and Nuñez–CyTOF the densest, and the strongest edge differed by context. The strongest edge overall, CD14*→*CD11B in Nuñez–CyTOF, reduced target uncertainty by 17.03% [95% CI 15.30, 18.91]. Some strong Nuñez–CyTOF edges linked markers of distinct lineages, such as CD19, CD21, and TCR*γδ*. These edges may reflect technical covariation, for example in staining intensity or channel crosstalk, rather than shared biology. Graph density should therefore not be read as biological connectivity.

**Figure 5.**
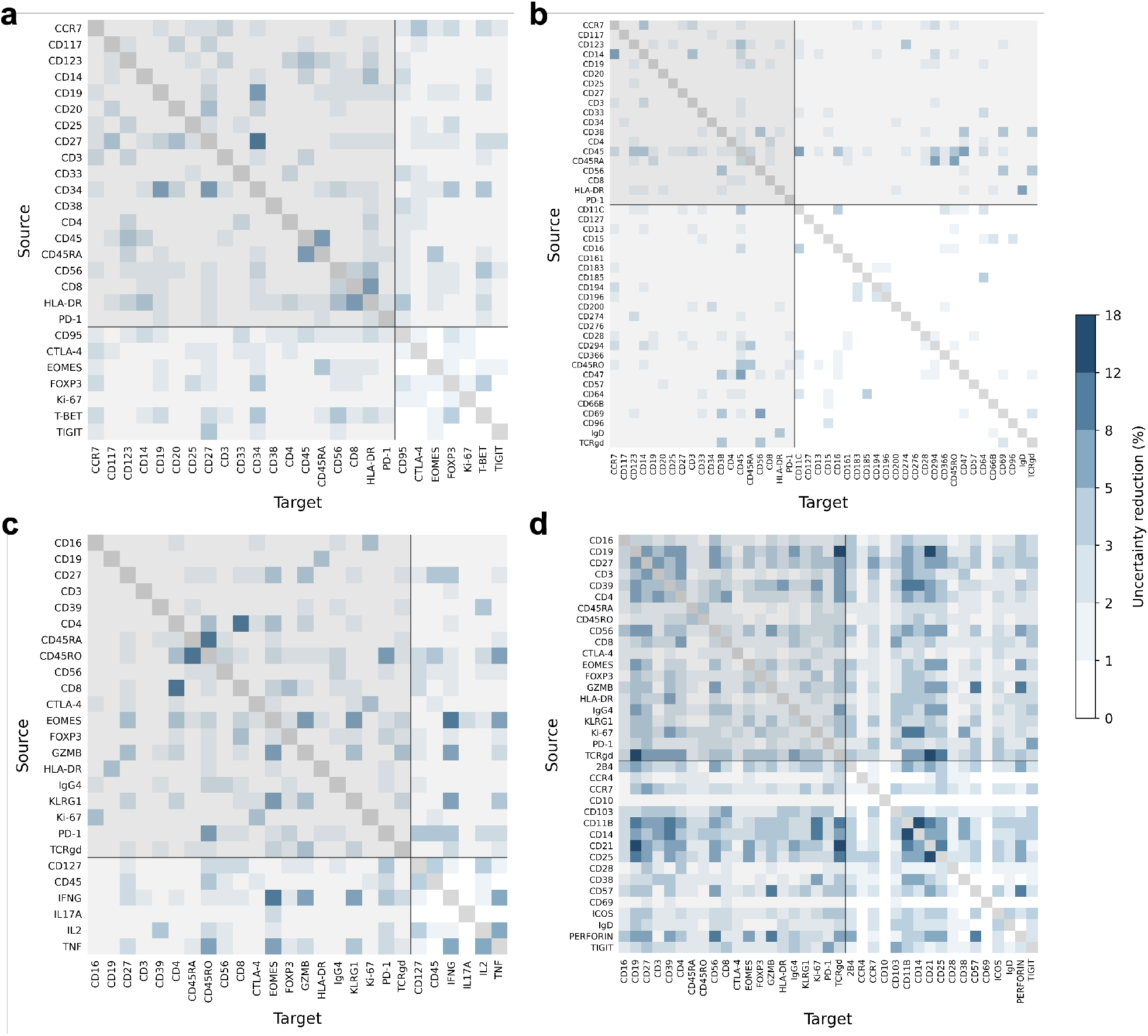
Cell-label-conditioned directional predictive-information atlas in four cytometry contexts: (a) AML–flow, (b) AML–CyTOF, (c) Nuñez–flow, and (d) Nuñez–CyTOF. Conditioning labels were manual-gating classes for AML and analysis-defined coarse lineage strata for Nuñez. Rows and columns denote source and target markers, respectively, and color shows the fraction of target uncertainty reduced by a stable edge. Gray diagonal cells are self-edges. White cells did not meet the stability criteria. Markers common to both modalities within each cohort are shown first, followed by modality-specific markers, with alphabetical ordering within each group. Light shading denotes common-marker rows and columns, and black lines separate common from specific markers. All panels use a shared stepped scale spanning 0–18%.

The Spearman correlation of common-edge magnitudes was below the prespecified transfer threshold of *ρ* = 0.60 in all three cross-context comparisons (Supplementary Table S7). Only AML– flow versus AML–CyTOF exceeded its marker-permutation null, supporting weak but nonrandom correspondence between the two graphs (*ρ* = 0.339, *q* = 0.009). Correlations were near zero between Nuñez–flow and Nuñez–CyTOF and between the two CyTOF contexts. The presence of some shared edges therefore did not imply transfer of the graph as a whole.

In nested held-out analyses of 133 target–context combinations, the second source added complementary information in 74.4%, both sources were informative but redundant in 15.8%, and a single source dominated in 6.0%. For example, combining perforin and CD57 in Nuñez–CyTOF reduced uncertainty about granzyme B by 14.83% [95% CI 13.43, 16.32], with an incremental gain of 3.24% [2.68, 3.86] from the second source. Most targets were therefore better explained by information distributed across multiple markers than by a single surrogate.

#### Graph-guided marker selection shows a trade-off between panel size and coverage

We varied panel size *K* to evaluate whether predictive-information graphs could guide marker selection. Reaching 95% of oracle pairwise coverage required 53.8% of markers (14/26) in AML–flow, 36.4% (16/44) in AML–CyTOF, 26.9% (7/26) in Nuñez–flow, and 21.6% (8/37) in Nuñez–CyTOF (Table 3; Figure 6). These panels reduced held-out MAE by 21.5–33.9% relative to median imputation and retained 77.8–87.6% of measured-full AML fine-label macro-AUPRC. The corresponding Nuñez coarse-stratum values were 36.9% and 40.9% and were treated as descriptive rule-consistency endpoints. By comparison, the common markers alone retained 89.7–100.6% of measured-full AUPRC for the same endpoints. The graph therefore quantified a trade-off between panel size and pairwise coverage. High coverage, however, did not imply that phenotype classification was preserved or that unselected markers could be dropped.

**Table 3.** Held-out utility of graph-guided panels required to reach 95% of oracle pairwise coverage. *K*_95_/full gives the number of selected markers and the number in the native full panel, and MAE reduction is relative to median imputation. Phenotype AUPRC retained is the ratio of macro-AUPRC for a classifier using only the *K*_95_ markers to that for the measured full panel. The endpoint was the manual-gating fine label for AML and the analysis-defined coarse lineage stratum for Nuñez; the latter is a descriptive rule-consistency endpoint. The *C*-only column, from the refit analysis in Supplementary Table S6, uses the 19 AML or 20 Nuñez common markers as a reference. MAE reduction is over the targets left unmeasured by each panel and is not directly comparable with Table 1.

| Context | $K_{95}/\text{full}$ | MAE reduction [95% CI] | AUPRC retained $K_{95}$ panel | AUPRC retained $C$ only |
| --- | --- | --- | --- | --- |
| AML-flow | 14/26 | 33.9% [31.2, 36.5] | 87.6% | 99.2% |
| AML-CyTOF | 16/44 | 21.5% [20.0, 23.1] | 77.8% | 89.7% |
| Nuñez-flow | 7/26 | 30.2% [28.2, 31.9] | 36.9% | 97.3% |
| Nuñez-CyTOF | 8/37 | 25.4% [24.3, 26.4] | 40.9% | 100.6% |

**Figure 6.**
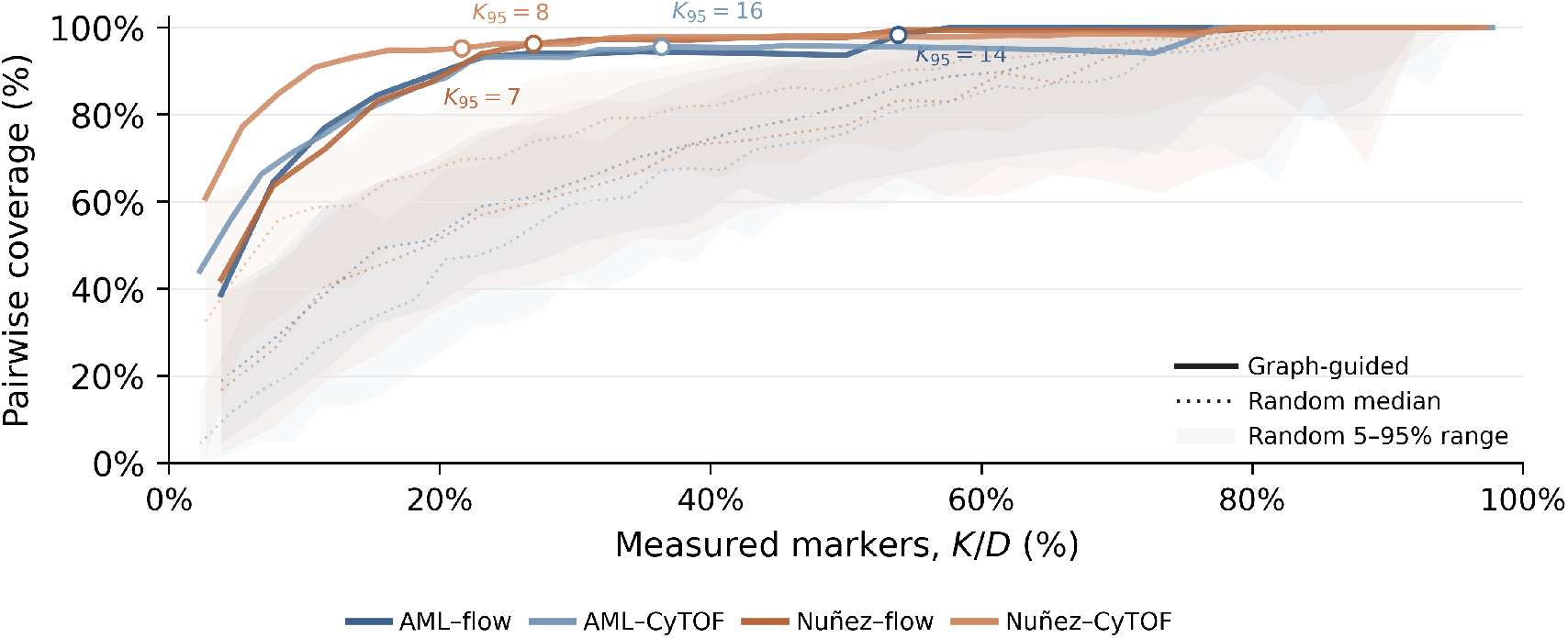
Oracle-relative pairwise coverage as a function of graph-guided marker-panel fraction. *K/D* is the number of selected markers divided by the native full-panel size in each context. Dark solid lines denote graph-guided panels. Light dotted lines and shading denote the median and 5th–95th percentiles of 100 random panels. Pairwise coverage credits each unmeasured target with its strongest single selected source. The oracle allows any other marker as that source. The labeled *K*_95_ is the first panel size to reach 95% of oracle coverage. Curves need not be monotone because the set of unmeasured targets changes with *K*. This metric is not a fraction of joint marker information.

Across these analyses, patient matching, source-exclusive markers, and within-platform marker relationships all carried information about target markers, but this information did not necessarily produce a net reconstruction or phenotype gain. Under the evaluated conditions, neither higher-capacity estimators nor graph-guided marker selection replaced information available through direct measurement.

## Discussion

In this study, how well unmeasured markers could be recovered in paired-platform cytometry depended on the information accessible to the model and on the resolution at which an output could be validated. Three stages were distinct: whether information existed, whether an estimator converted it into better reconstruction, and whether reconstructed markers contributed to downstream phenotypes. These stages diverged repeatedly across our analyses. The limits of cross-panel imputation should therefore be judged first by what the observed panel and pairing structure reveal about the target and by the resolution at which that information can be validated, not by estimator performance alone.

Within-platform masking showed that common markers could partially recover masked marker values, but marker-specific improvement and preservation of cell ordering varied widely. Median IQR retention was below 1 in every context and evaluable cell population. This contraction partly reflects median-based point prediction, which takes the median of neighboring values, and should not be read solely as a limit of the available information. Splitting a single full-panel dataset into pseudo-panels, or masking markers within one panel, has been used to provide same-cell truth for methods including CyTOFmerge(1), cyMAE(14), and UVAE(27). Because source and target measurements share an acquisition context, such experiments should be interpreted as controlled tests of conditional recoverability. Previous benchmarking has likewise shown that visually similar imputed distributions can coexist with substantial errors in cell-level expression and manual-gating results(23). Our paired cross-platform design extends this observation by separating the contributions of patient matching and source-exclusive inputs.

Across platforms, paired specimens conveyed sample context such as disease state and population composition, but their contribution depended on cohort and translation direction. Technical variation in cytometry includes instrument drift over time and batch effects that differ by marker, expression range, and cell population. This variation has motivated bead standards(9), multicenter harmonization(18; 30), and shared controls in normalization methods such as CytoNorm(33) and CytofRUV(32). Cross-platform comparisons in paired samples have also shown correspondence in broad lineages(11; 12), with platform-specific signal scales between CyTOF and flow cytometry(11). Our patient-matched bridge connects biological context and distributions but does not establish correspondence between individual cells from separate aliquots.

The source-exclusive permutation contrast further separated information access from usable reconstruction. *C* + *S* produced lower WD than *C* + *S*_perm_ in every direction, showing that the cell-level association between source-exclusive and common markers was informative for translation. However, *C* + *S* did not improve on *C*-only in any direction and was worse in both CyTOF*→*flow directions. Because within-specimen permutation also disrupts the joint structure of *S* and *C*, this contrast does not by itself establish information beyond *C*. In addition, adding shuffled source-exclusive markers increased WD relative to *C*-only in three directions (Supplementary Table S3), suggesting a cost of enlarging the input space. Early statistical file-matching work similarly noted that nearest-neighbor matching in common-marker space can create unobserved subpopulations(16).

Higher-capacity estimators did not consistently close the gap between available information and usable reconstruction. After a shared bridge, the best cross-platform model varied by translation direction. In the native-panel model sensitivity analysis, some nonlinear and generative models performed worse than median imputation in individual contexts. Shared-latent-space models such as CytoVI(13) and UVAE(27) provide flexible frameworks for heterogeneous-panel alignment, but latent alignment does not by itself guarantee patient-held-out reconstruction of target markers. These results do not show that kNN is universally superior to deep models, because each method’s native panel-integration pipeline and conditional sampling were not evaluated. Estimators should therefore be evaluated by marker-specific failures, distribution fidelity, and whether their information access is inductive or transductive, not by mean rank alone.

Biological utility was likewise distinct from reconstruction fidelity. For AML harmonized major phenotypes, both *C*-only and 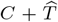 approached measured full panels, indicating that common markers already captured much of the information needed to distinguish broad lineages. In AML–CyTOF fine labels, however, measured full outperformed 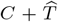 by 10.25 percentage points, and gaps exceeded 30 points for T cell *γδ* and T cell DN even after classifier refitting. Because 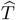 was computed from *C*, reconstruction could not add measurement information. These gaps instead identify phenotypes whose discrimination still depended on directly measured markers. Consistent with previous reports(1; 23), our results indicate that visual similarity or a single aggregate error should not be treated as evidence of biological equivalence. Utility must be validated for the intended downstream task.

Directional predictive-information graphs showed that marker information was not organized into a single universal network. Common-edge magnitude correlations were below the prespecified transfer threshold, and only AML–flow and AML–CyTOF showed weak preservation above the marker-permutation null. Reproducing a few familiar marker pairs therefore does not amount to transferring the overall predictive structure. In addition, most targets were better explained by combinations of markers than by a single surrogate. These edges are conditional associations that contribute to held-out prediction, not biochemical or causal interactions.

Graph-guided marker selection provided an exploratory framework for prioritizing markers under a constrained measurement budget. Existing panel-design methods score markers for discriminating target populations or seek compact gating strategies(6; 2), and CyTOFmerge proposed selecting informative shared markers for imputation(1). Our procedure instead optimizes held-out predictive relationships among markers. Panels reaching 95% of oracle pairwise coverage retained only 77.8–87.6% of measured-full AML fine-label macro-AUPRC. Two features of the objective may contribute to this gap. First, the graph was conditioned on cell labels, so information already explained by the labels did not count toward coverage. Second, coverage credited only the strongest single source for each target and ignored multi-source complementarity. Graph-guided panels should therefore narrow prospective panel candidates, not serve as universal minimal panels or replacements for direct measurement.

The principal limitations arise from the available data design. First, platform and panel changed together in the paired cross-platform measurements, so their effects could not be separated. Neither a within-modality multi-panel setting nor an umbrella panel measuring all source and target markers in one acquisition was available to define an empirical ceiling. Second, the cohorts were limited to AML and cancers treated with immune-checkpoint inhibitors(24). Because immune profiles depend strongly on population composition, our context-specific findings should not be read as universal properties of cytometry. The Nuñez strata were constructed from markers in *C*, the AML fine labels reflected a single institutional gating strategy defined from measured markers, and no sample-level clinical endpoint was evaluated. Third, estimator comparisons used point predictions under fixed training and preprocessing conditions. We did not evaluate conditional sampling, a source-blind empirical-distribution baseline, patient-level out-of-fold imputation of classifier training representations, label-free graph sensitivity, within-context graph reproducibility, or phenotype utility of same-size random panels. Future studies combining controlled multi-panel designs, umbrella references, multiple diagnoses, and independent annotations would further separate estimator, objective, and information limits.

For each intended use, Table 4 summarizes the information access and validation that our results suggest are required. In summary, paired specimens and marker imputation can connect sample composition and selected marker distributions across cytometry panels, but imputed values should not be assumed to provide cell-level information or phenotype resolution equivalent to direct measurement. Practical applications should jointly evaluate patient-held-out marker reconstruction, sample-level distribution fidelity, and downstream utility for every new platform and panel. Markers central to clinical or biological decisions should be measured directly. Predictive-information analysis can indicate which information a constrained panel may supplement and which remains unobserved.

**Table 4.**
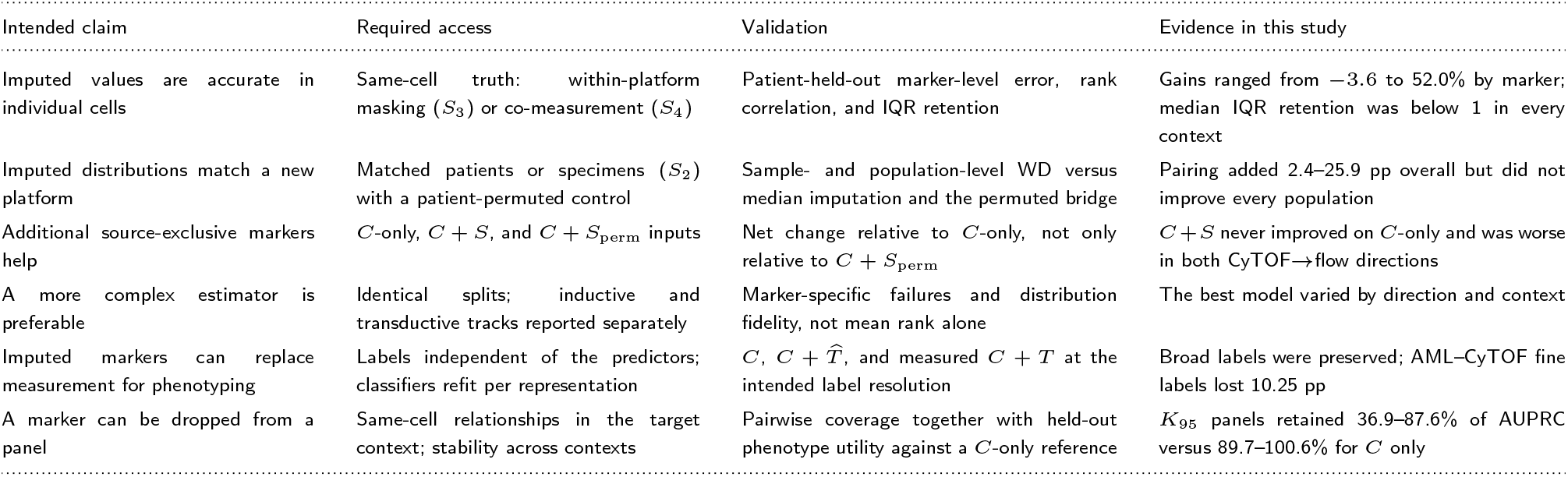
Practical checklist for evaluating cross-panel marker imputation. Each row pairs a downstream claim with the information access and validation it requires, together with the corresponding evidence from this study.

## Ethics statement

The I3H AML data were de-identified and analyzed in accordance with University of Pennsylvania Institutional Review Board requirements. The Nuñez dataset is a publicly released, de-identified cohort(24).

## Data availability

The I3H AML dataset is not publicly available because of institutional data-sharing restrictions. Access may be provided by the corresponding authors upon reasonable request, subject to institutional approval and applicable data-use agreements. The Nuñez dataset is publicly available from Zenodo at https://zenodo.org/records/5513853.

## Code availability

Code reproducing the Nuñez analyses from the public FCS files with exact kNN, including within-platform masking, patient-matched translation, phenotype classification, predictive-information graphs, and graph-guided panel selection, is available at https://github.com/jonghyunlee1993/CytoAlign_public. Source code for the Cytometry Relation Graph demonstration is available at https://github.com/jonghyunlee1993/cytometry_relation_graph. A hosted version of the interactive application is available at https://huggingface.co/spaces/jonghyunlee/cytometry-relation-graph.

## Author contributions statement

Methodology: J.L., A.N., C.-U.J., and M.I.; Resources: E.J.W., A.R.G., M.P.C., M.L.M., A.R., D.S., S.A.A., A.E.B., and S.; Supervision: D.K., E.J.W., and A.R.G.; Validation: J.J., M.E.L., L.T., Y.N., J.K., B.L., E.J.W., A.R.G., M.P.C., M.L.M., A.R., D.S., S.A.A., A.E.B., and S.; Writing – original draft: J.L. and D.K.

## Supplementary Methods

### Data Processing and Implementation Details

#### Data processing

AML data were exported as pre-gated, transformed events from the institutional pipeline and were not further transformed. Nuñez FACSymphony data were transformed as arcsinh(*x/*150), and Nuñez CyTOF data were used as released. Viability, barcode, and gate-source channels were excluded, and no additional quality-control gating was applied. Events were capped at 50,000 per specimen, resulting in 4.60 and 4.36 million cells for AML– flow and AML–CyTOF and 5.98 and 8.14 million cells for Nuñez–flow and Nuñez–CyTOF. Nuñez coarse strata were assigned from CD3, CD4, CD8, CD19, and CD56 using per-specimen thresholds at the intersection of a two-component Gaussian mixture, with the specimen median as fallback. One specimen required a fixed manual CD3 threshold because the mixture fit was unstable. For cross-platform population summaries, AML labels were harmonized to seven populations, with double-negative, double-positive, and *γδ* T cells grouped as Other T.

#### Splits and sampling

All analyses used five patient-grouped outer folds. Repeat visits from one patient were assigned to the same fold. Within-platform references sampled up to 5,000 cells per patient and 50,000 cells overall.

#### Cross-platform bridge

Common markers were mapped to patient-balanced percentile space using 101 quantiles fitted on outer-training cells. Balanced log-domain Sinkhorn transport (squared Euclidean cost divided by the number of common markers, *ε* = 0.05, 500 iterations) was solved between blocks of 1,000 source and target cells drawn from matched specimens, or from the deranged target patient’s specimens in the patient-permuted condition. Target-exclusive pseudo-targets were the barycentric projection of target values under the row-normalized transport plan. The resulting 50,000-cell bridge bank was balanced across patients and visits and was shared by all estimators in each fold.

#### Estimators

kNN used *k* = 50 with Euclidean distance and the neighbor median. Ridge regression used *α* = 1. Gradient-boosted trees used histogram gradient boosting per target (learning rate 0.08, 200 iterations, 31 leaves, L2 penalty 1). The MLP had two hidden layers of 128 GELU units and was trained with AdamW (learning rate 10^*−*3^, weight decay 10^*−*5^), smooth-L1 loss, batch size 4,096, up to 30 epochs, and early stopping with patience 5 on a held-out validation set grouped by patient within platforms and by bridge-bank specimen across platforms. CytoVI (scvi-tools 1.5.0.post1) used a 10-dimensional latent space, one 128-unit hidden layer, a mixture prior, up to 100 epochs, and batch size 1,024. UVAE (commit 37bf8a5) used a 50-dimensional latent space, two 256-unit hidden layers, 20 epochs, and batch size 512.

#### Phenotype classifiers

Classifiers were class-balanced multinomial logistic regressions on standardized features (inverse regularization 1, L-BFGS, 1,000 iterations) trained on up to 10,000 cells per class. For *C* + *T*, training cells were imputed by kNN excluding the cell itself, and held-out cells by standard kNN. The same classifier was used for the *K*_95_ panels.

### Scaling and Reconstruction Metrics

Marker values were standardized using outer-training patient-balanced locations and robust scales. The primary point metric for within-platform masking was relative MAE reduction versus the source-blind median,

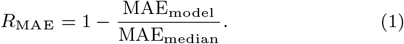

A value of 0 indicates performance equal to median imputation, and a positive value indicates lower error. Same-cell Spearman correlation evaluated preservation of cell ordering.

The primary distribution metric was WD, defined as the 1-Wasserstein distance for each held-out specimen and target marker divided by the robust outer-training target scale. Secondary metrics were normalized errors in the median, IQR, 90th percentile, and positive fraction. IQR retention was the predicted IQR divided by the observed IQR. The positivity threshold was the intersection of a two-component Gaussian mixture fitted in the outer-training data. When this threshold could not be estimated reliably, the outer-training median was used.

### Cross-platform Contrasts

Within each outer fold, target-exclusive pseudo-targets *T* were generated by an entropic optimal-transport bridge in patient-balanced percentile space. Target markers, cell-type labels, and endpoint labels were not used to fit the bridge. The patient-matched bridge was compared with a bridge fitted after permuting patient keys. For Table 2, the patient-first WD of each target marker (specimens averaged within patient, then patients averaged) was expressed relative to that marker’s median-imputation WD, and improvements were averaged across markers with equal weight. The patient-permuted condition was averaged over five fixed patient derangements. Confidence intervals for the pairing contribution were obtained from 2,000 patient-bootstrap replicates that recomputed the full marker-averaged statistic. Figure 3a instead reports the ratio of direction-level mean WDs, which weights high-WD markers such as CD57 and CD103 more heavily. Source-exclusive input was assessed by comparing *C*-only, *C* + *S*_perm_, and *C* + *S*. Each contrast was defined as WD under the condition on the left minus WD under the condition on the right.

### Phenotype Classification and Statistical Inference

Classifiers were refit separately for each representation using outer-training patients and evaluated in held-out patients. The 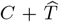 training representation was generated by leave-self kNN imputation of outer-training cells. The primary contrast subtracted measured-full (*C* + *T* ) AUPRC from *C*-only or 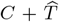 AUPRC. Reconstruction gain was the difference in macro-AUPRC between 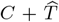 and *C*-only. Applying the measured-full classifier directly to other representations was reported separately as a classifier-shift sensitivity analysis.

Results from specimens, and from the three MLP seeds in within-platform analyses, were first averaged within patient. Other analyses used a single seed. Confidence intervals were calculated with 2,000 bootstrap replicates using patients as the resampling unit. Marker and population heterogeneity in cross-platform translation was tested with 5,000 within-patient label permutations. Multiple testing was controlled with the Benjamini–Hochberg procedure.

### Predictive-information Graph Estimation

Graphs were estimated in the four native full-panel contexts using patient-held-out five-fold cross-validation. Within each outer fold, marker-specific empirical percentiles were estimated from training cells. To prevent tied values from being divided among bins, expression was discretized into eight states using the midpoint of the left and right empirical ranks. The weight for training cell *j* was

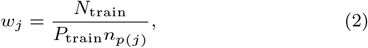

so that each patient had equal total weight. Here, *N*_train_ and *P*_train_ are the numbers of training cells and patients, respectively, and *n*_*p*(*j*)_ is the number of cells contributed by the patient containing cell *j*.

Conditional probabilities for the target state were estimated from weighted frequency tables with Dirichlet smoothing (*α* = 0.5):

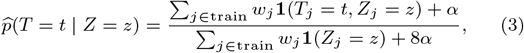

where the conditioning variable *Z* was the cell label *L* in the baseline model and (*L, M* ) in the augmented model. Here, *L* denoted the manual-gating label in AML and the analysis-defined coarse lineage stratum in Nuñez. For held-out cell *i*, the log-score gain from adding source marker *M* was

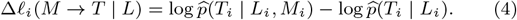

Under the true conditional probability, the expectation of this quantity is the conditional mutual information *I*(*T* ; *M* | *L*). We used held-out prediction and blocked permutation to reduce finite-sample bias and apparent associations induced by patient-specific marker distributions. DREMI(15) renormalizes the conditional density so that values across the source range are weighted equally when characterizing the functional strength of a relationship. In contrast, our estimate is weighted by the observed cell distribution, evaluated on held-out patients, and conditioned on cell labels. It therefore reflects predictive utility for imputation rather than signaling influence.

Each marker was permuted independently within patient*×*label blocks, preserving marginal distributions within patient and label while removing same-cell marker correspondence. Primary edge information for patient *p* was the observed gain minus the mean gain across 20 permuted graphs:

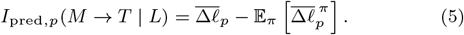

Edges were reported in nats per cell and as the target predictive-information fraction

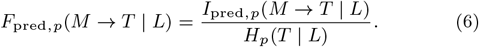

Here, *H*_*p*_(*T* | *L*) is the held-out cross-entropy of the baseline model and represents target uncertainty remaining after prediction from the label alone. Thus, *F*_pred_ is the fraction of that uncertainty reduced by the source marker.

For patient-level edge estimates, 95% confidence intervals were calculated with 2,000 bootstrap replicates, and positive information gain was tested with a one-sided patient-level *t*-test. Benjamini– Hochberg correction was applied within each context and graph layer. An edge was stable if its confidence-interval lower bound exceeded 0, *q ≤* 0.05, and its mean information fraction was at least 1%.

Preservation across contexts was evaluated for three prespecified pairs (the two within-cohort cross-platform pairs and the cross-cohort CyTOF pair), using every directed edge among markers common to both contexts and harmonized major-cell labels. The primary metric was the Spearman correlation of common-edge magnitudes (information fractions). Secondary metrics were the stable-edge Jaccard index and node-coverage rank correlation. A null distribution was generated by 1,000 permutations of marker names within one context, and *ρ ≥* 0.60 was prespecified as a practical transfer threshold.

Multi-source structure for each target was evaluated in a nested outer-fold analysis. Within each outer-training fold, the two leading sources *M*_(1)_ and *M*_(2)_ were selected by information fraction after subtracting the mean of three blocked permutations. Null-corrected gains were then re-estimated in held-out patients using 20 permutations. Additional information from the second source was

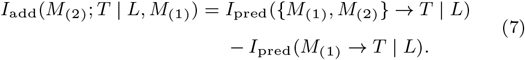

An individual source was supported if the lower bound of its 95% bootstrap confidence interval exceeded 0 and its information fraction was at least 1%. Incremental information additionally required a one-sided Benjamini–Hochberg-adjusted *q ≤* 0.05. A relationship was classified as *multi-source complementary* when both *M*_(1)_ and *I*_add_ were supported, *pairwise redundant* when both individual sources were supported but *I*_add_ was not, *single dominant* when only *M*_(1)_ was supported, and *unsupported* otherwise.

### Graph-guided Marker Selection

The information-saturation sweep was performed on the aggregate graph for each context. Edges with information fractions below 1% were set to 0. Coverage supplied by panel *P* to targets not yet measured was

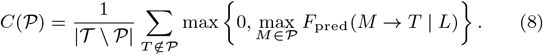

For each unmeasured target, coverage was normalized by an oracle denominator that allowed every other marker as the single strongest source. Oracle-relative pairwise coverage was the ratio of *C*(*P*) to this oracle, summed over the same unmeasured targets. Because both terms use the strongest single source, this ratio does not measure joint information from multiple sources, and the target set changes with *K*. Starting from an empty panel, markers were added sequentially to maximize the increase in *C*(*P*). With *D* denoting native full-panel size, every *K* through *D −* 1 was evaluated. Sweeps were compared across contexts by measured-marker fraction *K/D*, and we identified the smallest *K* that first reached 95% of oracle pairwise coverage. The comparator comprised 100 random panels of the same size.

Held-out validation was performed at *K*_95_. Panels were reselected within each outer-training fold. Selected markers were scaled by outer-training medians and IQRs, patient-balanced references were capped at 50,000 cells, and exact kNN (*k* = 50) was applied. Marker-wise medians among neighboring target values were used as predictions, and relative MAE reduction was calculated against the outer-training target median. Phenotype utility compared a multinomial logistic classifier using only selected markers with a measured-full-panel classifier, using AML fine labels and Nuñez coarse strata as distinct endpoints. All confidence intervals were calculated with 2,000 patient-bootstrap replicates. Because *K* itself was selected post hoc from the aggregate graph, this analysis was interpreted as an exploratory sensitivity analysis.

### Interactive Software Demonstration

We provide an interactive Cytometry Relation Graph demonstration for exploring the predictive-information framework in independent cytometry datasets. The application accepts preprocessed patient-level cytometry files measured with a common marker panel, estimates a label-free directional predictive-information graph, and displays a relation heatmap, incoming and outgoing information for each marker, marker-set pairwise coverage, an interactive network, and connected-component structure. Its built-in example uses a 10-patient subset of the public Nuñez CyTOF dataset.

The demonstration is available at https://huggingface.co/spaces/jonghyunlee/cytometry-relation-graph, and source code is available at https://github.com/jonghyunlee1993/cytometry_relation_graph. The application differs from the manuscript analysis in three ways. It does not use cell labels, it uses within-patient rather than patient*×*label blocked permutation with fewer permutation and bootstrap replicates, and it reports exact minimum-cardinality marker sets rather than the greedy *K*_95_. It is therefore intended for exploring the framework in new data rather than for reproducing the label-conditioned predictive-information atlas in the manuscript. Only public or non-sensitive data should be uploaded to the public Space. Private clinical or controlled-access data should be analyzed with a local deployment.

**Figure S1.**
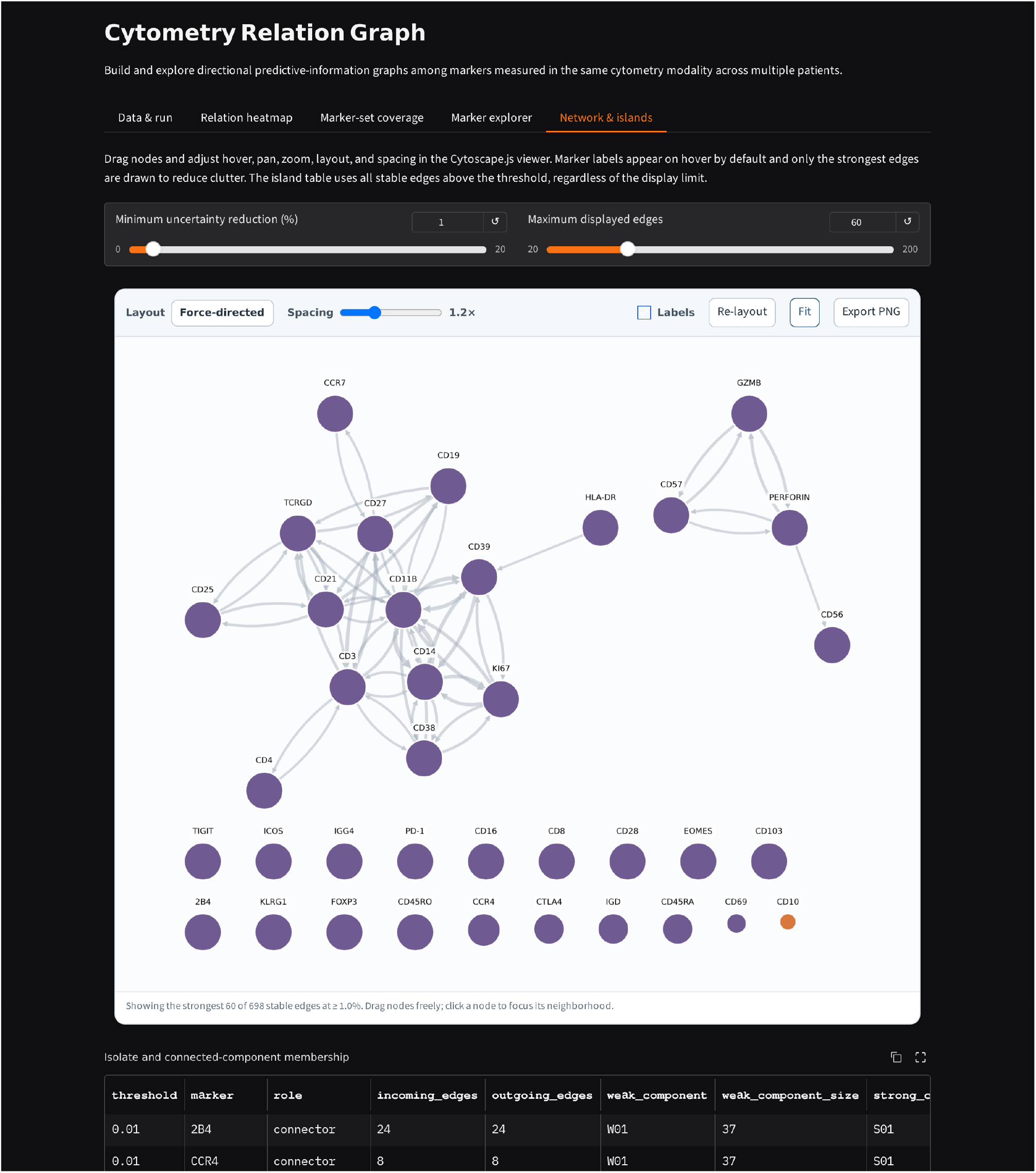
Interactive Cytometry Relation Graph demonstration. The screenshot shows the *Network & islands* view for the built-in public Nuñez CyTOF example. Users can adjust the minimum uncertainty-reduction threshold and number of displayed edges and explore the directional marker network, isolated markers, and connected-component membership. Separate tabs provide the relation heatmap, marker-set pairwise coverage, and marker explorer. Estimates from this label-free demonstration do not reproduce the label-conditioned predictive-information atlas in the manuscript.

**Table S1.**
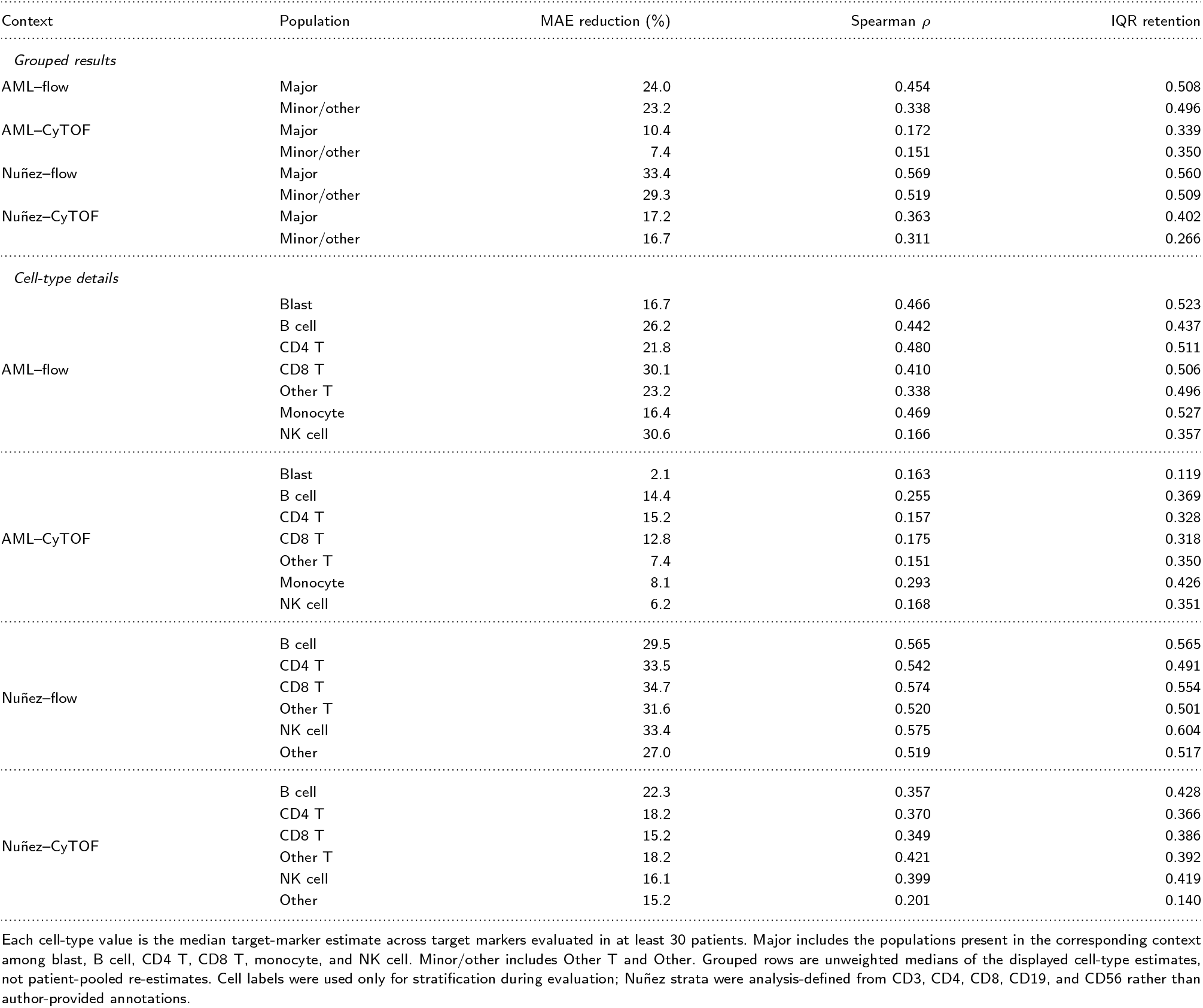
Grouped and cell-type-specific kNN same-cell reconstruction.

## Supplementary Results

### Cell-type Reconstruction Details

Cell-type-specific MAE reduction ranged from 2.1% for Blast in AML–CyTOF to 34.7% for CD8 T cells in Nuñez–flow. IQR retention was below 1 for every cell type (0.119–0.604), showing that distributional contraction was not confined to a particular population (Table S1).

### Cross-platform Translation Details

Supplementary Table S2 reports marker-level WD for patient-matched *C*-only translation. In AML–flow*→*AML–CyTOF, the three lowest-WD markers were CD64 (0.241), CD200 (0.299), and CD11C (0.302), whereas the three highest were CD57 (6.055), CD161 (2.328), and CD66B (1.380). In AML–CyTOF*→*AML–flow, CD95 (0.330), CTLA-4 (0.392), and TIGIT (0.503) had the lowest WD, whereas EOMES (0.720), T-BET (0.658), and Ki-67 (0.577) had the highest. In Nuñez–flow*→*Nuñez–CyTOF, CD10 (0.325), 2B4 (0.406), and CCR4 (0.422) had the lowest WD, whereas CD103 (63.232), IgD (1.284), and TIGIT (1.130) had the highest. In Nuñez–CyTOF*→*Nuñez–flow, CD45 (0.247), IL17A (0.319), and TNF (0.406) had the lowest WD, whereas IFNG (0.456), IL2 (0.423), and CD127 (0.413) had the highest.

**Table S2.** Marker-level WD for patient-matched *C*-only translation.

| Translation context | Median [IQR] | Lowest WD | Highest WD |
| --- | --- | --- | --- |
| AML–flow→AML–CyTOF | 0.651 [0.546–0.851] | CD64 (0.241) | CD57 (6.055) |
| AML–CyTOF→AML–flow | 0.538 [0.447–0.617] | CD95 (0.330) | EOMES (0.720) |
| Nuñez–flow→Nuñez–CyTOF | 0.654 [0.525–0.826] | CD10 (0.325) | CD103 (63.232) |
| Nuñez–CyTOF→Nuñez–flow | 0.409 [0.341–0.420] | CD45 (0.247) | IFNG (0.456) |

**Table S3.**
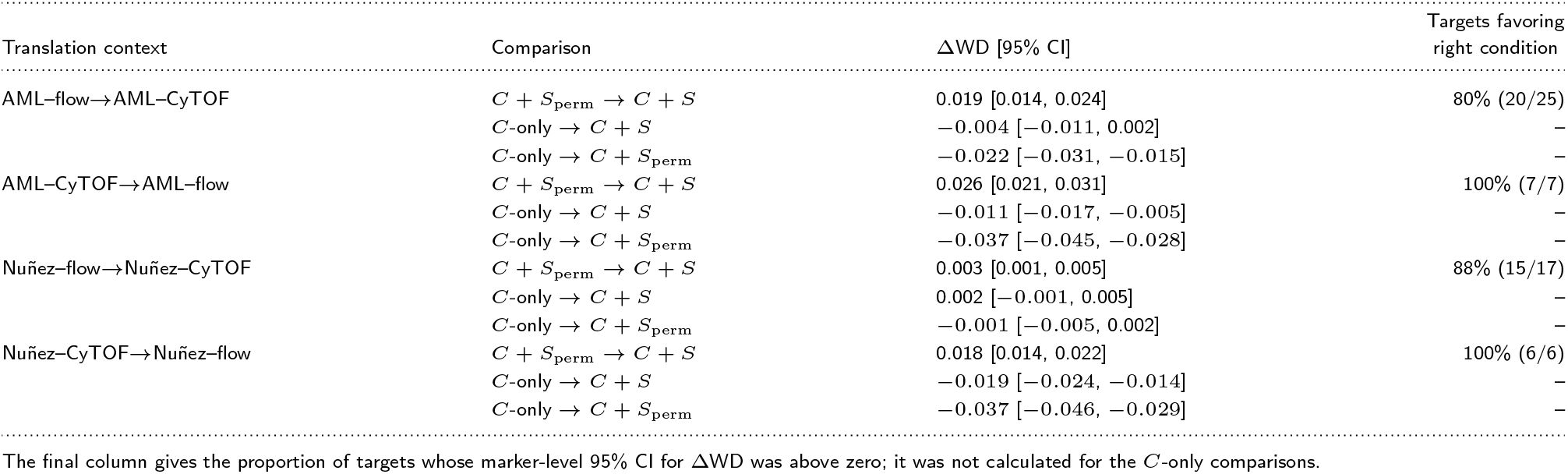
Cross-platform WD contrasts from source-exclusive inputs. ΔWD is the WD under the condition to the left of the arrow minus that under the condition to the right; positive values indicate lower WD for the right-hand condition. The *C*-only *→ C* + *S*_perm_ contrast measures the cost of adding shuffled source-exclusive dimensions.

*C* + *S* produced lower WD than *C* + *S*_perm_ in all four directions. However, the WD reduction for *C* + *S* relative to *C*-only did not have a positive 95% confidence interval in any direction, and adding shuffled source-exclusive markers increased WD relative to *C*-only in three directions.

**Table S4.**
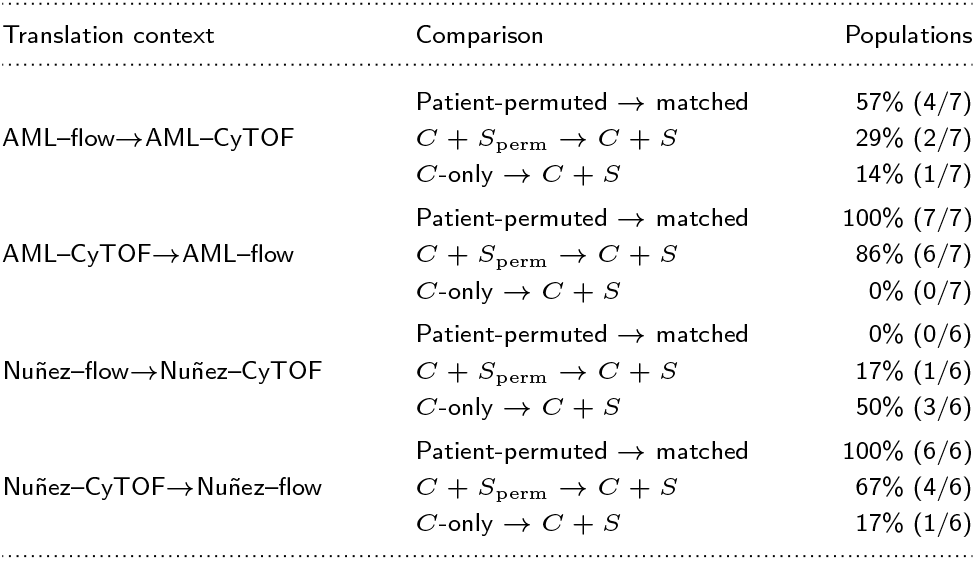
Proportion of harmonized populations with lower WD under the condition to the right of the arrow in cross-platform comparisons. Parentheses give the number of such populations and the total number evaluated: seven in AML, including Other T (double-negative, double-positive, and *γδ* T cells), and six in Nuñez. Nuñez populations were analysis-defined coarse lineage strata rather than author-provided annotations.

Population-level results are shown in Supplementary Table S4. Patient-matched translation produced lower WD than patient-permuted translation in every major population for both CyTOF-to-flow directions but in no population for Nuñez–flow*→*Nuñez–CyTOF. The two source-exclusive input contrasts also varied by population.

Marker and population heterogeneity was significant in every translation direction (Benjamini–Hochberg-adjusted *q <* 0.001). In Nuñez–flow*→*Nuñez–CyTOF, sparse CD103 produced a marker mean of 4.349 despite a median of 0.654.

### Native-panel Model Sensitivity

Native-panel model sensitivity results are shown in Supplementary Table S5.

**Table S5.**
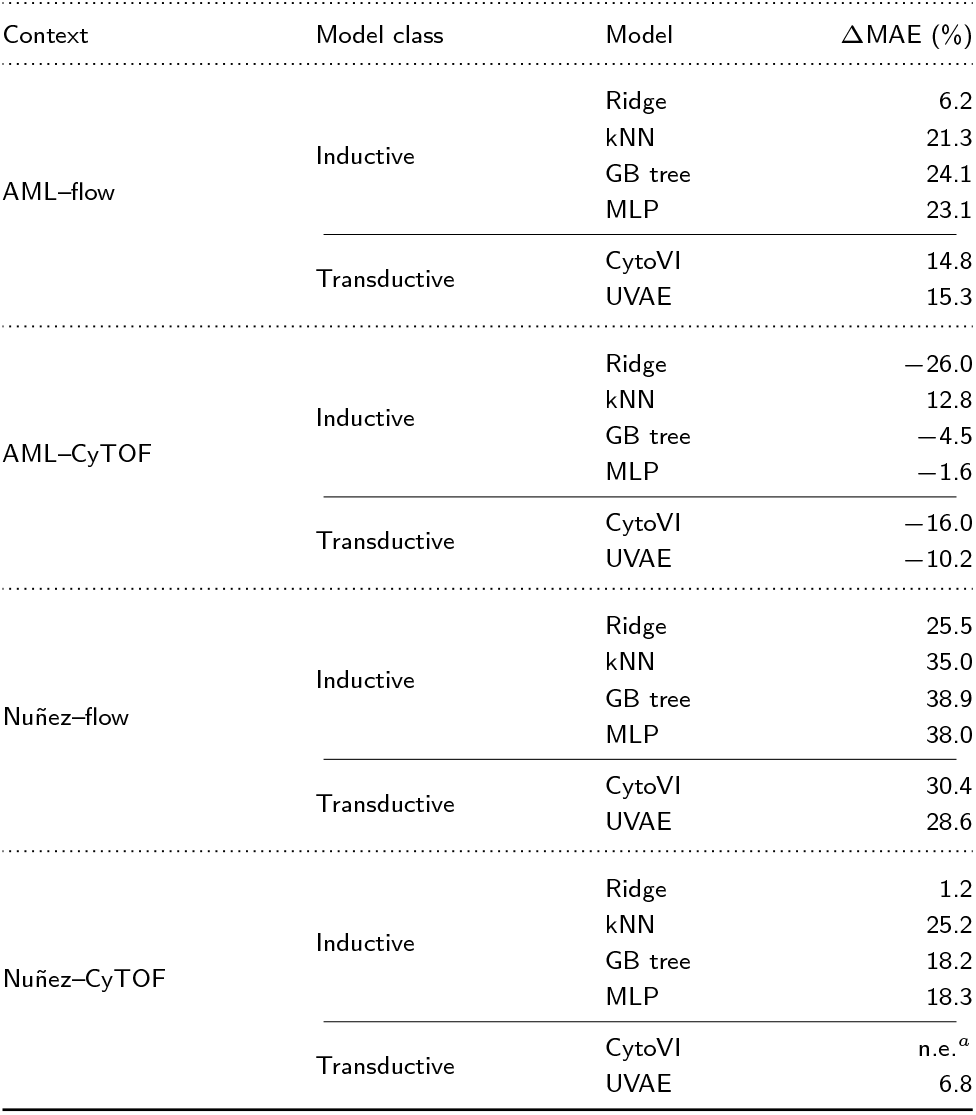
Native-panel model sensitivity. Values are MAE reductions (%) relative to median imputation; positive values indicate lower error. Inductive and transductive models were evaluated in separate prespecified pipelines and were therefore not ranked directly across model classes. kNN, CytoVI, and UVAE values use the robust marker scaling described in Methods; ridge regression, gradient-boosted trees (GB tree), and the MLP standardize or bin inputs internally and are invariant to this scaling. ^*a*^Not estimable: CytoVI training diverged or produced extreme predictions in at least one fold under both marker-scaling rules; the previously reported value came from a run in which no fold diverged.

### Phenotype-classification Refit Sensitivity

With the same 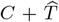 input, classifier refitting increased AML–CyTOF fine-label AUPRC by 4.5 percentage points relative to the fixed classifier (Table S6). This comparison isolates classifier– representation shift but does not separately identify the effect of distributional contraction.

**Table S6.** Phenotype-classification contrasts and sensitivity to classifier refitting. Measured full is patient-first mean AUPRC; the remaining columns report percentage-point differences with 95% patient-bootstrap confidence intervals. Negative values in the first two contrasts indicate lower performance than the measured full panel, whereas positive values in the final contrast indicate improvement from refitting. Only analysis-defined coarse lineage strata were reported for the Nuñez cohort; these are descriptive rule-consistency endpoints rather than independent phenotype annotations.

| Context | Labels | Measured full | $C$-only — full<br>95% CI | Refit $C + \hat{T}$ — full<br>95% CI | Refit — fixed<br>95% CI |
| --- | --- | --- | --- | --- | --- |
| AML-flow | Major | 0.985 | -0.9 [-1.3, -0.7] | -0.9 [-1.3, -0.6] | +0.3 [+0.2, +0.4] |
| AML-flow | Fine | 0.982 | -0.8 [-1.1, -0.5] | -0.8 [-1.1, -0.5] | +0.4 [+0.2, +0.6] |
| AML-CyTOF | Major | 0.979 | -3.0 [-3.5, -2.5] | -3.1 [-3.6, -2.5] | +1.2 [+0.9, +1.5] |
| AML-CyTOF | Fine | 0.973 | -10.1 [-10.9, -9.3] | -10.3 [-11.1, -9.5] | +4.5 [+3.7, +5.3] |
| Nuñez-flow | Major | 0.820 | -2.2 [-2.5, -1.9] | +1.3 [+0.9, +1.6] | +0.9 [+0.6, +1.2] |
| Nuñez-CyTOF | Major | 0.880 | +0.5 [+0.4, +0.7] | +0.2 [+0.1, +0.3] | 0.0 [-0.1, 0.0] |

### Predictive-information Graph Details

Cross-context graph comparisons are shown in Supplementary Table S7.

**Table S7.**
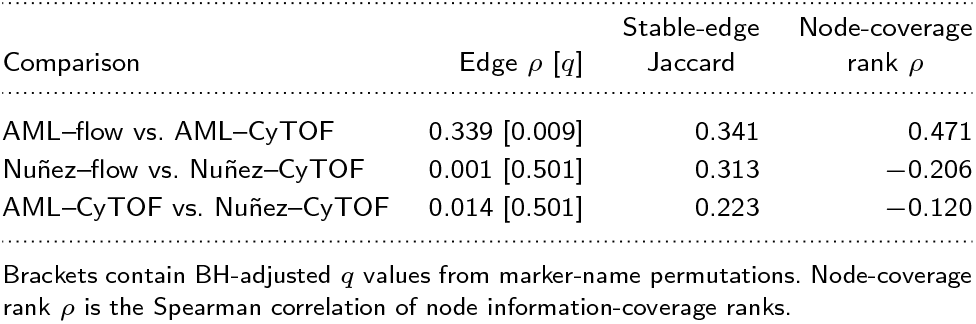
Preservation of predictive-information graphs across contexts. The primary comparison is the Spearman correlation of shared directional-edge magnitudes.

Context differences were also evident in the integrated within-label atlas (Figure 5). Nuñez–CyTOF had higher proportions of stable and strong edges than Nuñez–flow, but these quantities do not represent recovery rates for directly measured values or causal connectivity. Edge counts and the strongest edge in each context are reported in Supplementary Table S8.

The strongest edge in each context reduced target uncertainty by 7.47–17.03%. Large directional edges were observed in every context, but the marker pairs differed, so no single relationship can be interpreted as a platform-independent biological rule.

**Table S8.**
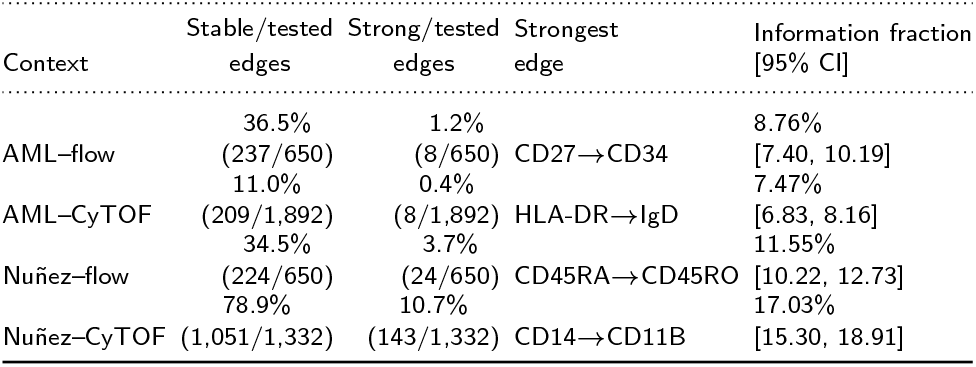
Predictive-information graph summary by context. The denominators for stable- and strong-edge proportions are the tested directional edges; strong edges are stable edges with an information fraction of at least 5%.

